# Alteration of mechanical stresses in the murine brain by age and hemorrhagic stroke

**DOI:** 10.1101/2023.09.25.559368

**Authors:** Siyi Zheng, Rohin Banerji, Rob LeBourdais, Sue Zhang, Eric DuBois, Timothy O’Shea, Hadi T. Nia

## Abstract

Residual mechanical stresses in tissues arise during rapid differential growth or remodeling such as in morphogenesis and cancer. These residual stresses, also known as solid stresses, are distinct from fluid pressures and dissipate in most healthy adult organs as the rate of growth decreases. However, studies have shown that residual stresses remain substantially high even in mature, healthy brains. The genesis and consequences of these mechanical stresses in a healthy brain, and in aging and disease remain to be explored. Here, we utilized and validated our previously developed method to map residual mechanical stresses in the brains of mice in three different age groups: 5-7 days, 8-12 weeks, and 22 months old. We found that residual solid stress increases rapidly from 5-7 days to 8-12 weeks in mice, and remains high even in mature 22-month-old mice brains. Three-dimensional mapping of the residual stresses revealed an increasing trend from anterior to posterior in coronal sections of the brain. Since the brain is rich in negatively charged hyaluronic acid, we evaluated the contribution of charged extracellular matrix (ECM) constituents in maintaining solid stress levels. We found that lower ionic strength leads to elevated solid stresses, a finding consistent with the unshielding effect of low ionic strength and the subsequent expansion of charged ECM components. Lastly, we demonstrated that hemorrhagic stroke, accompanied by loss of cellular density, resulted in decreased levels of residual stress in the murine brain. Our findings contribute to a better understanding of the spatiotemporal alteration of residual solid stresses in healthy and diseased brains, a crucial step toward uncovering the biological and immunological consequences of this understudied mechanical phenotype in the brain.

**Significance Statement:** While emerging evidence highlights the importance of solid stresses in embryogenesis and tumor growth, the genesis and consequences of residual solid stresses in the adult normal brain remain poorly understood. Understanding the spatiotemporal distribution and alteration of the residual solid stresses as the brain ages and is impacted by neuropathologies, such as a stroke, will elucidate the biological and immunological consequences of maintaining these stresses. This study suggests solid stress could serve as a potential biomarker in aging and diseases associated to the brain.

## Introduction

Residual solid stresses are the compressive and tensile mechanical stresses generated and transmitted by solid components of tissue as opposed to interstitial fluid pressure (1). Differential growth, cell-cell and cell-matrix interactions, and electroosmotic stresses involved in organ development and disease progression give rise to residual mechanical stresses (2–5). Residual solid stresses in biological tissues were first identified in arteries, where changes in systemic pressure can trigger asymmetrical growth in the inner and outer layers (6), and later also found during rapid growth, a defining feature of organogenesis (3, 5–8) and tumor growth (1, 9–11). During tumor growth, residual solid stresses were shown to be a key contributor in tumor progression (12, 13), immune evasion (14), and treatment resistance by directly compressing the blood and lymphatic vessels (11, 15, 16) or directly compressing the cancer cells (17). More recently, we showed that solid stresses applied on the neurons surrounding brain tumors causes neuronal death (18, 19), which can be protected by neuroprotective strategies such as the use of lithium (18). While elasticity, a traditionally studied biomechanical property, has been extensively probed in brain development and pathogenesis (20–23), our understanding of the genesis and consequences of residual solid stress is limited in brain.

When the growth and remodeling of tissues are slowed down, as in most adult organs, solid stresses appear to dissipate as previously shown in liver and kidneys (10); in contrast, solid stresses in the adult brain are maintained at high levels. The first evidence on existence of solid stresses in adult brain was shown by the partial cut method developed by Y.C. Fung (24). In a later work, it was shown that the differential growth in brains, synthetically modeled by polymer expansion, is responsible for the brain folding (25). In a recent study where the temporal volumetric change of different brain anatomical compartments were quantified over the lifespan of over 100,000 humans (26), differential volumetric changes of key components of the brain was reported: (i) gray matter reached its maximum volume in childhood (1-3 years) while white matter reached its maximum volume in young adulthood (20-30 years), and (ii) while the gray matter, white matter, and cortical volumes decline in late adulthood (60-100 years), the ventricle volume dramatically increases during this time. The differential volumetric changes imply the existence of residual solid stresses in the human brain. The brain also undergoes substantial morphological and structural alterations in diseases such as Alzheimer’s (22, 27, 28), stroke (29–31), scar formation (20), and cancer (18, 19) that potentially give rise to residual mechanical stresses. Key unanswered questions are how the residual solid stress changes in these diseases, and whether these mechanical stresses contribute to the progression of the disease. Our limited understanding of the solid stress-associated mechanobiology in the brain is partially due to challenges in directly mapping these stresses. Unlike stiffness, a mechanical properties which has been successfully demonstrated as a key player in the brain health and disease (20–23), measuring the solid stress requires either mechanical intervention to relax the existing stresses (10, 18, 32) or embedding microgels to estimate the stresses (33–37).

Here, we use our previously developed method (10, 32) to map the solid stresses in a mouse brain at three different time points during its lifespan: early childhood (5-7 days), adulthood (8-12 weeks), and late adulthood (22 months). Our method, based on relaxing solid stresses by slicing the freshly resected brain into thin sections, provides the deformation induced by stress relaxation, which is quantified as (i) the normalized deformation, (ii) average local curvature, and (iii) changes in surface area. We then studied viscoelastic properties of the brain at these age groups through an unconfined compression test to obtain the Young’s modulus, rate-dependent stiffening, and stress relaxation time constant of the brain and their alteration by age. To better understand the determinants of solid stresses in brain, we then investigated the effect of ionic strength on the residual solid stress in the mouse brain and found that lower ionic strength results in higher level of solid stress, suggesting that charged ECM constituents may contribute to the maintenance of solid stress in brain. We evaluated the 3-D distribution of the residual solid stress in a series of coronal slices along the sagittal plane, and found that solid stresses increase from anterior to posterior sections. Finally, we studied brains with a hemorrhagic stroke and found that regions with stroke have lower levels of residual solid stress. The unique approach and conceptual findings of this study will contribute to a better understanding of the origin and biological consequence of solid stress, a potential biomarker in brain aging and pathologies.

## Results

### Residual solid stress in the murine brain increased with age

Based on the first principle that tissues containing residual solid stress deform upon release of physical confinement (10, 32), we sliced brains taken from mice ranging from 5-7 days to 22 months old to quantify the alteration of the residual solid stress as the brain matures. The harvested brain was embedded in agarose prior to slicing after which the 250 µm thick tissue slice was left in phosphate buffered saline without the agarose surrounding so the residual solid stresses relax (**Fig. 1A, B**). The tissue slice, deformed after stress relaxation, was embedded in agarose and fixed with formalin to preserve the deformed structure for 3-D fluorescent microscopy (**Fig. 1C, D**). The structural changes imaged are quantified as the normalized deformation (**Fig. 1E**), changes in surface area (**Fig. 1F**), and local curvature (**Fig. 1G**). To validate our measurement method and rule out the effect of buoyancy in the measurement, we quantified and compared the solid stress indices, including normalized deformation, area ratio, and mean curvature, before and after flipping the slice and found no significant difference between the groups across all indices (**Fig. S1**). These results demonstrated that the buoyancy had no significant effect on deformation and solid stress quantification, and thus did not cause any artifact in our analysis. Next, we compared the solid stress indices in freshly excised vs fixed slices of the same tissue and found no significant difference in the deformation and solid stress quantification (**Fig. S2**).

**Figure 1.**
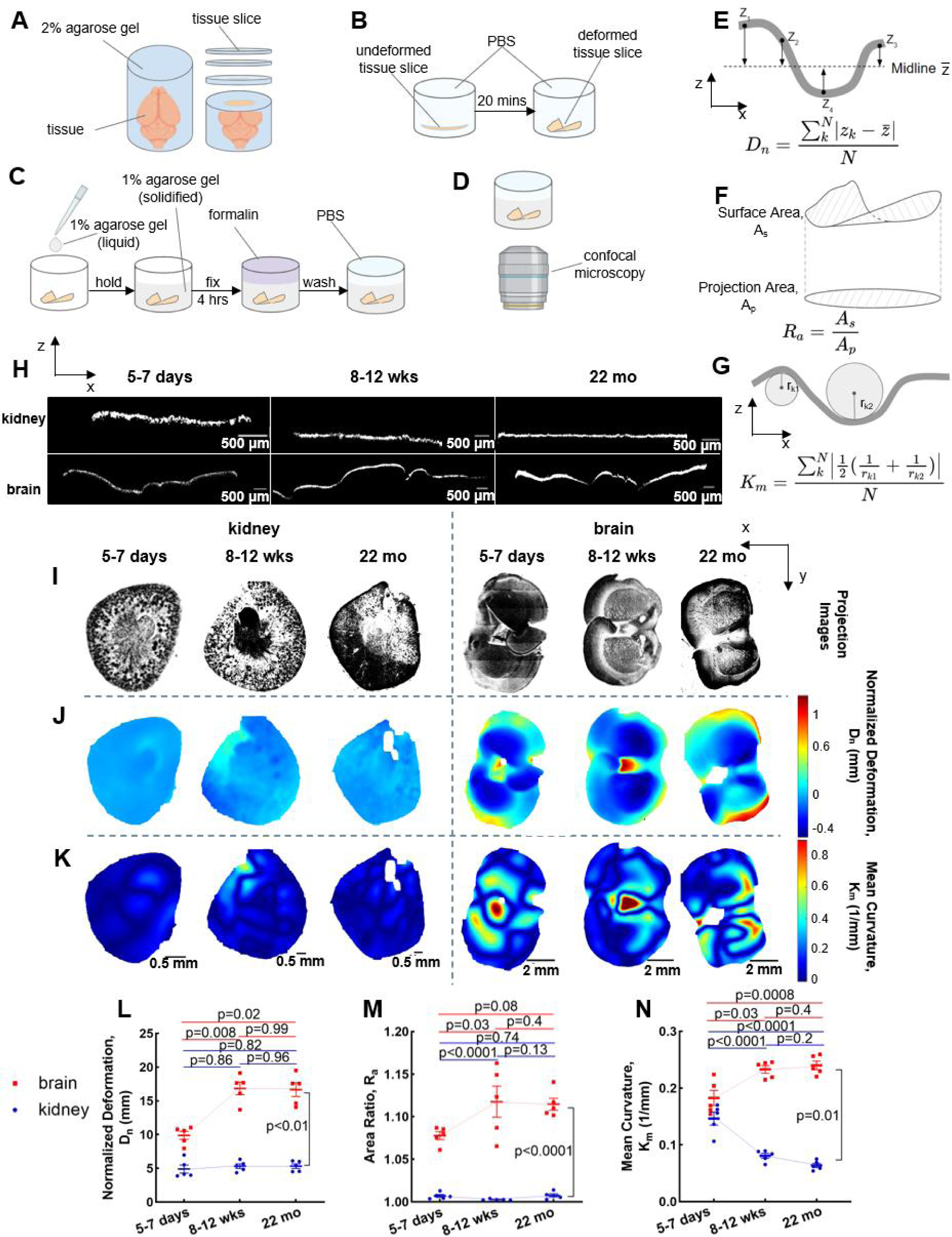
Residual solid stresses in murine brain are age-dependent. (**A**) The fresh brain is embedded in 2% agarose, then sliced with a compresstome to obtain a specific thickness of 250 μm. (**B**) Slices are left in a phosphate buffered saline at room temperature for 20 minutes to deform as the solid stress is released in the tissue slice. (**C**) The deformed slices are embedded in 1% agarose, then fixed with formalin overnight, and washed with PBS. (**D**) The deformed slice is imaged via confocal microscopy. (**E**) The normalized deformation, Dn, an index of residual solid stress, is defined as the average height difference from curved surface to the midline. (**F**) The area ratio, Ra, defined as the ratio of surface area and projection area, is used as another index of solid stress. (**G**) The curvature, k, defined as the reciprocal of the curvature radius, is used as the third index of solid stress. Mean curvature, Km, is the average of mean curvature on tissue slices. (**H**) The orthogonal views of deformation of representative brain and kidney (negative control) slices from 5-7 days, 8-12 weeks, and 22 months mice. (**I**) The projection images, (**J**) deformation maps, and (**K**) mean curvature maps of representative brain and kidney slices from 5-7 days, 8-12 weeks, and 22 months mice. (**L**) Residual solid stresses are estimated in multiple ways by quantifying the normalized deformation, (**M**) area ratio, and (**N**) mean curvature. Brain slices have significantly higher normalized deformation, area ratio, and mean curvature than kidney slices in all age groups. Brain slices have increased normalized deformation, area ratio, and mean curvature from 5-7 days to 8-12weeks, and then keep plateau from 8-12 weeks to 22 months (mean ± SEM, N=5 mice, two-tailed t-test).

We compared the deformation of the brain slices against kidney (known to have negligible residual solid stress (10)) slices which remained flat with no local bending (**Fig. 1H**) taken from the same mouse. Representative deformation map of kidney slices from microscopy in each age group (**Fig. 1I**) showed a relatively flat surface and the deformation was negligible and uniform across the cut surface indicating lower residual solid stresses (**Fig. 1J**). Compared to this, the deformation maps of brain slices show substantially larger and unevenly distributed deformations indicating higher residual solid stresses (**Fig. 1J**). Similarly, the curvature maps of representative kidney and brain slices showed higher curvature unevenly distributed in brain slices, in contrast to flat kidney slices (**Fig. 1K**). The normalized deformation and area ratios across each age group for all experimental repeats was significantly higher for brain slices than the kidney as the negative control (**Fig. 1L, M**). The mean curvature, and the brain slices also had significant higher curvature than kidney slices (**Fig. 1N**) other than when the tissue was taken from 5-7 days old mice. Overall, the normalized deformation, area ratio and mean curvature corroborate that the brain had a larger residual solid stress than the kidney. We also studied the heart as a positive control, as it is known to have high residual solid stress (8), and found that the heart slices showed greater and unevenly distributed deformation compared to kidney (**Fig. S3**).

Next, we evaluated the changes in the residual solid stress in brain with age. We found that the normalized deformation in brain slices increased significantly between the 5-7 day and 8-12 week age groups, and then remained constant between the 8-12 week to 22 month age groups (**Fig. 1L**). We observed consistent trend in other solid stress indices, the area ratio and mean curvature (**Fig. 1M, N**). We expected the fast prenatal growth to give rise to higher growth-induced stresses, which would dissipate in slower postnatal brain development. We were surprised to find that the peak stress in the brain occurs long after birth up till late adulthood (22 months). Unlike other organs (e.g., kidney and liver) (10), the solid stress in healthy adult brains do not dissipate over time, and high levels of stress are maintained even into late adulthood. While we are aware of the major differences between the human and mouse brain, the differential change of volume of different brain components in humans (**Fig. S4**) can be informative in understanding the age-related dynamics of solid stress. Unfortunately, due to the invasive nature of our solid stress measurement technique, it is not possible to transform the lifespan geometrical changes reported for humans (26) to more functional mechanical readouts such as solid stresses. To better understand the genesis and age-related dynamics of solid stress in the brain and how it is maintained we require a systematic and longitudinal analysis of the mouse brain as whole and its separate components (26).

### Age-dependence of viscoelastic properties in murine brain

Next, we asked whether the age-dependent changes in solid stress is associated with age-dependent alteration of materials properties of the brain. We quantified the equilibrium and instantaneous Young’s modulus, rate-dependent stiffening, and stress relaxation time constants of these properties in normal brain in the same age groups (**Fig. S5**). Unlike solid stress indices, equilibrium Young’s modulus significantly increased between the 5-7 day and 8-12 wee age groups, and then decreased between the 8-12 week to 22 month age groups (**Fig. 2A**). While the stiffening between 5-7 day to 8-12 week age groups is consistent with an increase in solid stress in this period, the softening of the brain that happens in 22 month old mice trends opposite to the elevated solid stress in this age group. The rate-dependent stiffening, measured as the ratio of instantaneous to equilibrium modulus and an indicator of brain viscoelasticity, showed only a decreasing trend (p = 0.06) from 5-7 days mice to 22 months mice (**Fig. 2B**). We reported the stress relaxation time constant in terms of fast and slow time constants, representing the early and late parts of the stress relaxation curves. The fast relaxation time constant decreased significantly from the 5-7 day to 8-12 week age groups, and remained low for the 22 month age group, while the slow relaxation time constant remained unchanged across age groups (**Fig. 2D**). This showed that mice in early infancy have longer time-dependent viscoelastic behavior compared to young and late adulthood. While the stress relaxation curve cannot be used to fully decouple intrinsic viscoelasticity (flow-independent) from poroelastic response (flow-dependent), the fast relaxation time constant is often associated to poroelastic behavior(39), which implies that the brain in early infancy has lower hydraulic permeability compared to later in life.

**Figure 2.**
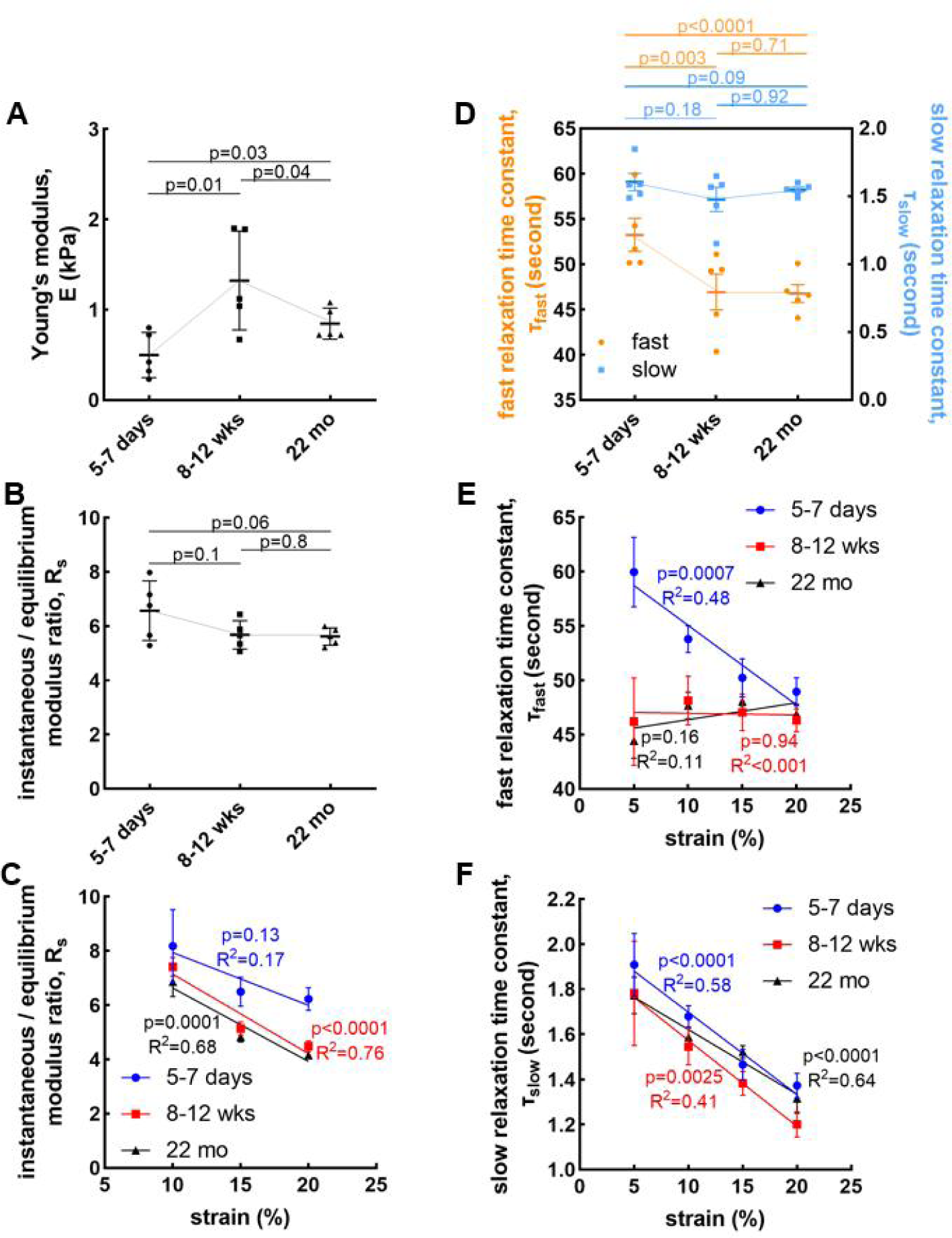
Viscoelastic properties in murine brain are age-dependent. (**A**) Young’s modulus and (**B**) rate-dependent stiffening defined as instantaneous / equilibrium modulus ratio are age-dependent in brain (mean ± SEM, N=5 mice, two-tailed t-test). Brain from 8-12 weeks mice has the highest Young’s modulus among all age groups. Instantaneous / equilibrium modulus ratio has a decreasing tendency in mice brains from 5-7 days to 22 months. (**C**) The instantaneous / equilibrium modulus ratio is strain-dependent (mean ± SEM, N=5 mice, one-way ANOVA test). Brain tissue from 8-12 weeks and 22 months mice have decreasing tendency of instantaneous / equilibrium modulus ratio with strain. (**D**) Fast relaxation time constant, and slow relaxation time constant, changes with age (mean ± SEM, N=5 mice, two-tailed t-test). Brain from 5-7 days mice has both the highest fast and slow relaxation time constant. The changing tendency of (**E**) fast and (**F**) slow relaxation time constant with strain (mean ± SEM, N=5 mice, one-way ANOVA test).

Since the relaxation of solid stresses results in large deformations (**Fig. 1**), we next probed whether the mechanical properties of the brain are linear or strain dependent. We observed that the rate-dependent stiffening, and the relaxation time constants are all strain-dependent in all the age groups (**Fig. 2E, C, and F**) except the fast relaxation time constant seen in the 8-12 week and 22 month groups. While the data demonstrates the highly nonlinear material properties of the brain across age groups, the underlying mechanisms need further investigation of the brain constituents and their response at high strains. Interestingly however, this nonlinear behavior remains mostly unchanged from infancy to late adulthood (**Fig. 2E, C, and F**).

### Residual solid stresses are higher in anterior than posterior sections of the murine brain

To better understand the spatial variation of the solid stress in brain, we probed the variation of solid stress from the anterior to posterior of the brain in 8-12 weeks old mice. The normalized deformation maps showed larger deformations moving towards the posterior sections (**Fig. 3A**). The curvature maps also showed larger and unevenly curvature existed on brain slices with the moving more posteriorly (**Fig. 3B**). The quantification of normalized deformation and area ratio had lower values in the first multiple slices, starting from the posterior sections, and higher values in the final slices with a statistically significant increasing trend (**Fig. 3A**). Interestingly the data showed a local minimum at about 750 μm and a local maximum at about 3 mm from the front of the brain. These local minimum and maximum appeared consistently in all the three solid stress indices used in this study, the normalized deformation, mean curvature, and area ratio. It should be mentioned that since all these metrics are normalized by the area of the slice, the changes in the area of the slices from anterior to posterior sections are ruled out in these readouts, and do not present any confounding effect.

**Figure 3.**
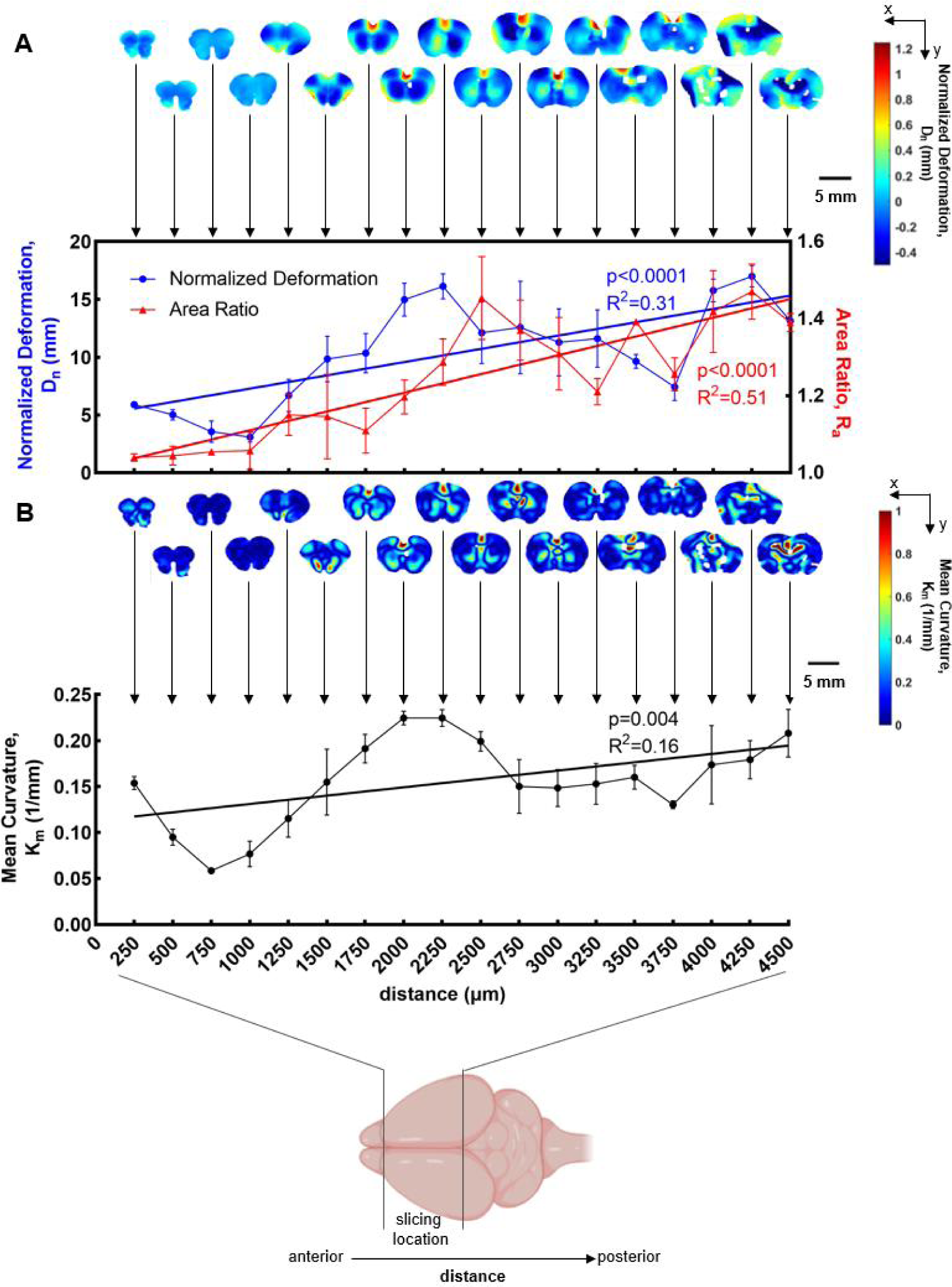
Residual solid stress distribution in the whole brain. (**A**) The normalized deformation and area ratio of brain slices at a coronal direction (mean ± SEM, N=3 mice, one-way ANOVA test) demonstrated a consistent increasing trend from the anterior to posterior slices. (**B**) The mean curvature of brain slices at a coronal direction (mean ± SEM, N=3 mice, one-way ANOVA test) demonstrates an increasing trend from the anterior to posterior sections.

Brain extracellular matrix (ECM) components contribute to structural and functional properties of the brain and play a significant role in brain development. Previous research showed that ECM proteins which are especially expressed in mature brain, mainly aggrecan, brevican, and tenascin-R, are heterogeneously distributed throughout the brain (40). Such unique and complex distribution of ECM plays an important role in brain physiology and might affect the solid stress distribution. The result of our study that higher residual solid stresses exist in posterior mature brain would have a potential relationship with the ECM distribution and be helpful to understand the connection between the mechanical properties and biological components of the brain.

### Residual solid stress in the murine brain depends on ionic strength

There is an abundance of negatively-charged extracellular matrix constituents in brain such as hyaluronic acid, which can contribute to maintenance of solid stresses as in tumors (41) and connective tissues (39, 42). These negatively-charged glycosaminoglycans (GAGs) repel each other and induce a local electro-osmotic expansion, which can contribute to the observed high level of solid stresses in the brain. We hypothesized that when the ionic strength is decreased, e.g., by decreasing the salt concentration, the charged constituents are further unshielded which increases the effect of expansion forces that ultimately increases solid stress. We measured the solid stress in 8-12 weeks old mice, in different ionic strength of 0.01X, 0.1X, 1X, and 10X PBS. Representative confocal microscopy images, deformation maps and curvature maps from four tissue slices taken from a similar region of the brain are subjected to different salt concentrations (**Fig. 4A-C**). At low ionic strength of 0.01X PBS, the brain slices from the similar region of the brain showed larger deformation and larger curvature indicating substantially higher levels of solid stresses. We observed that all the solid stress indices were substantially increased by decreasing the ionic strength (**Fig. 4D-F**). Unexpectedly, we did not see any significant reduction in normalized deformation, area ratio, and mean curvature when the ionic strength was increased from 1X to 10X. This key observation implies that the electro-osmosis associated with the charged constituents of the brain (e.g., GAGs) may contribute to the high level of solid stresses. This observation has important implications in diseases where the GAG content in brain is altered, such as mucopolysaccharidosis disorders, Alzheimer’s disease, schizophrenia, Parkinson’s disease, epilepsy, and brain tumors (43–46).

**Figure 4.**
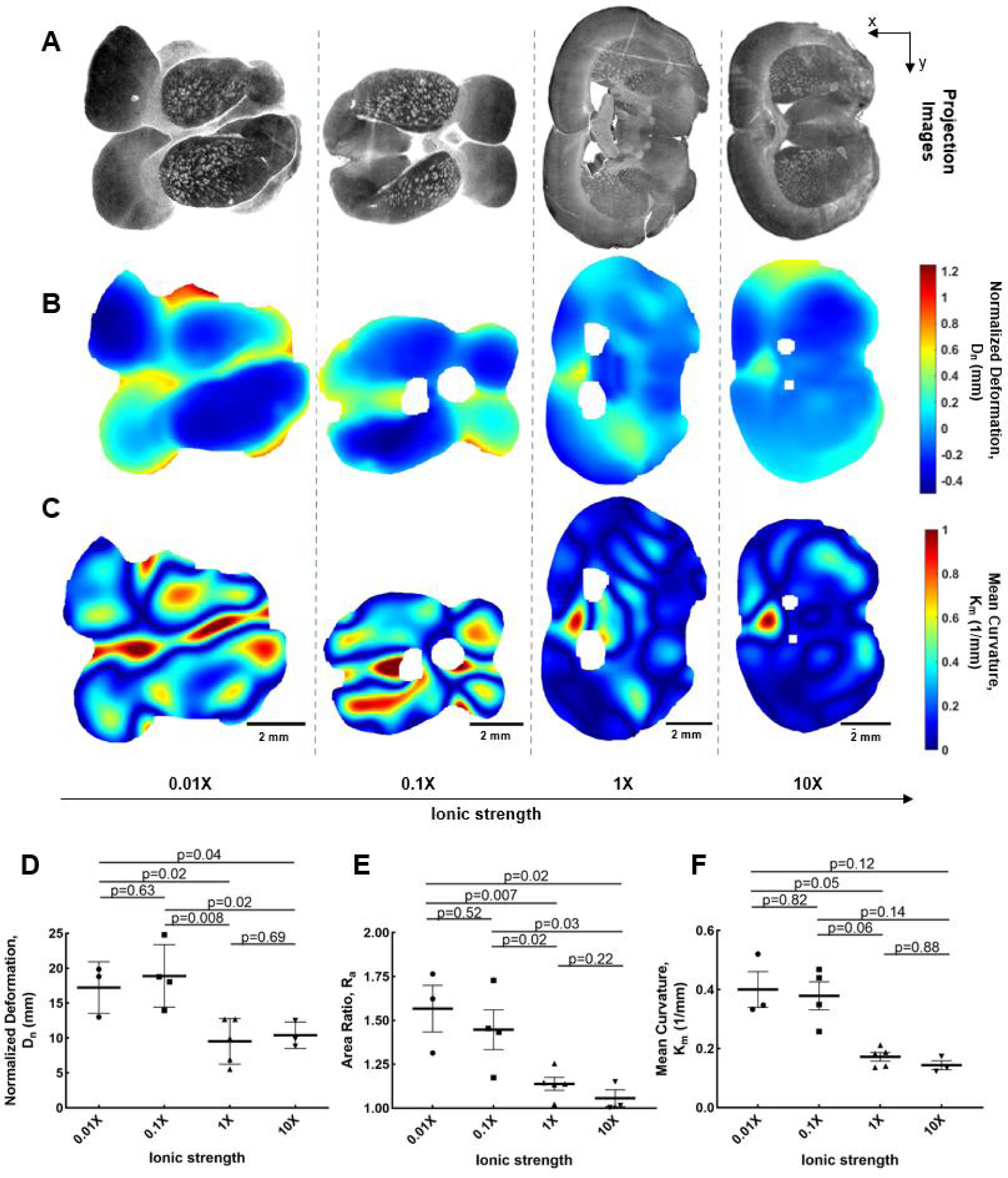
The effect of ionic strength on residual solid stresses. (**A**) Projected microscopy images, (**B**) normalized deformation maps and (**C**) mean curvature maps of representative brain slices from 8-12 weeks mice under different ionic strength. (**D**) The normalized deformation (**E**) area ratio and (**F**) mean curvature shows a decreasing trend with increasing ionic strength (mean ± SEM, n=3-5 slices, two-tailed t-test).

### Hemorrhagic stroke results in decreased residual solid stress in murine brains

Finally, we decided to probe how CNS injury such as hemorrhagic stroke affects the solid stress levels in brain. Hemorrhagic stroke causes a massive and irreversible change in ECM composition and morphology. Initial disruption of local vessels results in the efflux of blood from the damaged vessels, formation of a hematoma and subsequent edema along with the creation of a cytotoxic microenvironment leading to the death of local neural cells (47–49). The death of local neural cells results in the infiltration of monocytes, such as neutrophils and subsequently, and the recruitment of peripheral macrophages (50). The infiltrated, Cd13-positive immune cells phagocytose cellular debris and in conjunction with locally-recruited, Gfap-positive astrocytes, take part in the formation of the astroglial border, which effectively walls off the region of tissue damage from the preserved, viable CNS tissue (**Fig. 5A, B**). In addition to the loss of cellular density within the lesion core, there is also a depletion of NeuN-positive neurons on the periphery of the lesion core (**Fig. 5A**). Due to these structural and cellular alteration in hemorrhagic stroke, we hypothesized that solid stress levels will substantially alter compared to the normal brain. Analysis of the hemorrhagic stroke using the slicing method indeed revealed a loss of symmetry within the brain, as evidenced by gross morphology (**Fig. 5C**), as well as deformation (**Fig. 5D**) and curvature (**Fig. 5E**) mapping of coronal brain sections. Further quantification of the normalized deformation, area ratio, and mean curvature revealed that brains with hemorrhagic strokes had a significant decrease in all three metrics (**Fig. 5F-H**), demonstrating that hemorrhagic stroke significantly reduced residual solid stress in murine brains. This observation demonstrates that solid stress can be used as a mechanical biomarker which is sensitive to cellular and structural changes in the brain. Furthermore, the reduction of solid stresses in the hemorrhagic stroke lesion may contribute to the pathophysiology of stroke due to mechano-responsiveness of immune and brain cells. Such mechanical cues may lead to identification of new biomarkers and therapeutic targets in future studies.

**Figure 5.**
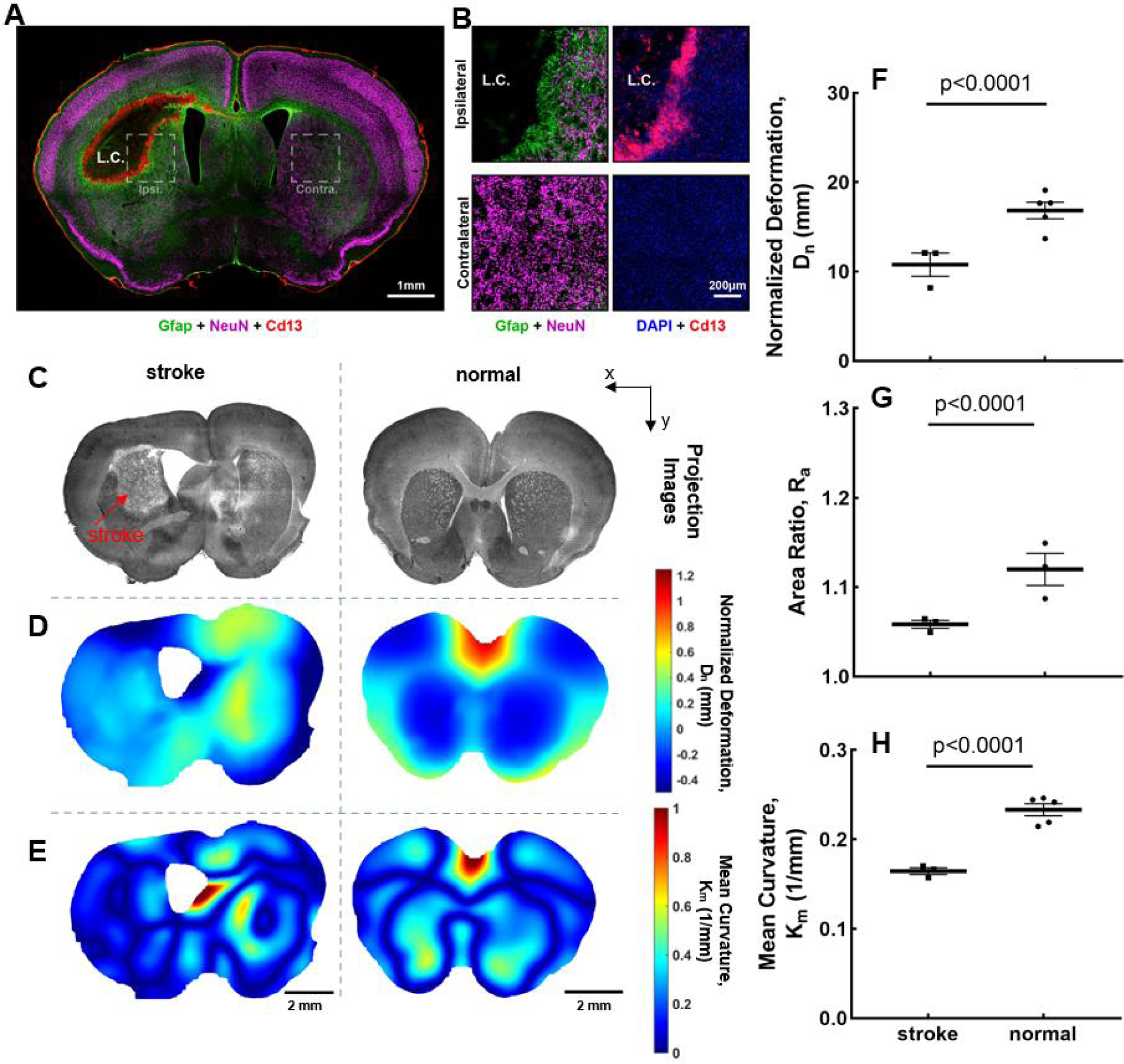
Hemorrhagic stroke results in lower residual solid stress in murine brains. (**A**) The loss of cellular density within the lesion core, and the depletion of NeuN-positive neurons on the periphery of the lesion core. (**B**) The infiltrated, Cd13-positive immune cells phagocytose cellular debris and in conjunction with locally-recruited, Gfap-positive astrocytes, take part in the formation of the astroglial border. (**C**)The projected image from microscopy, (**D**) deformation maps, and (**E**) mean curvature maps of representative brain sections from hemorrhagic stroke and healthy mice. (**F**) The normalized deformation, (**G**) area ratio, and (**H**) mean curvature are lower in lobes with hemorrhagic stroke compared to normal brain sections (mean ± SEM, N=3 mice, two-tailed t-test). Healthy brain sections have higher normalized deformation, area ratio, and mean curvature than hemorrhagic stroke brain sections.

## Discussion

In this study, we quantified and mapped residual solid stress in murine brains at three different stages during the mouse lifespan at 5-7 days, 8-12 weeks, and 22 months. We utilized our previously developed tissue slicing method (10, 32) to quantify the deformation induced by relaxation of the intrinsic solid stress. We quantified three separate indices of solid stress -- normalized deformation, area ratio, and mean curvature -- and validated our method against both a positive- and negative-control. We found that lower ionic strength solutions increase levels of solid stress in the murine brain. We evaluated the distribution of residual solid stress up to 5 mm into the brain by making serial coronal tissue slices and found that higher solid stresses exist in posterior sections than in anterior sections in mature murine brain. At last, we showed that solid stress is lowered in case of a hemorrhagic stroke in the mature murine brain. The findings of this study will help to better understand the role of solid stress as a potential biomarker for aging and different brain pathologies.

One of the limitations of this study is the invasiveness of our approach to measure solid stress, which method cannot be used in human brains in vivo. To demonstrate solid stress as a sensitive biomarker, a noninvasive method is required to probe the longitudinal changes in solid stress in healthy and diseased brains. Potential solutions are the recently developed methods where a sensor of solid stress, e.g., deformable microgels, are embedded in the organ of interest (33–37). While these methods are either minimally invasive during the injection of the sensor (37), the sensor can potentially be embedded during the early development as in the case of tumor growth (33). Developing a method to non-invasively study solid stresses in healthy brain of animals and human remains an unmet need with immense potential for uncovering the role of mechanical forces in brain development and aging.

Another unmet need is the in vitro and in vivo models of solid stress in brain. We previously developed in vivo models to apply chronic compressive stresses on brain (18, 19) to recapitulate the compressive forces applied by tumors onto the surrounding brain tissue. However, a model system that is able to faithfully recapitulate the normal growth-induced solid stresses in brain is still an unmet need. Such in vitro or in vivo models can substantially improve our understanding of mechanobiology and mechano-immunity associated with solid stress in brain development, aging and disease.

## Materials and Methods

### Mice Strain

Mice were housed and bred under pathogen–free conditions at the Boston University Animal Science Center. All experiments conformed to ethical principles and guidelines under protocols approved by the Boston University Institutional Animal Care and Use Committee (BU IACUC). All mice were C57BL/6 strain from three different age groups: 5-7 days, 8-12 weeks, and 22 months. A breeding pair of transgenic B6.129(Cg)-Gt(ROSA)26Sortm4(ACTB-tdTomato,-EGFP)Luo/J (JAX #007676, Jackson lab, ME), hereafter referred to as mTmG, was purchased to start a colony and was the primary source of all animals for the experiments. mTmG mice were used in the 5-7 day and 8-12 week age groups and endogenously expressed tdTomato fluorescence in all cell membranes. Mice in the 22 month age group were wild type. N=5 mice were used for all age groups.

### Slicing method to measure residual solid stresses

#### Releasing residual solid stress

We used our previously developed tissue slicing method (10). Briefly described here, 2% agarose is prepared using low gelation temperature agarose (Sigma-Aldrich) mixed with phosphate-buffered saline (PBS). The 2% agarose solution is in a liquid state at a temperature of 40°C. The organs, including brain and kidney, were immediately extracted post-euthanasia through CO_2_, and embedded in 2% agarose in a stainless-steel cast provided by the commercial vibrating microtome, Compresstome (F00395A, Precisionary Instruments LLC). The agarose-embedded organs are fully immersed in PBS and sliced via the Compresstome to the desired thickness of 250 μm which is thin enough for showing the deformation, but thick enough to avoid the tissue being broken by handling (**Fig. 1A**). The tissue slice is detached from the agarose slice spontaneously or manually with a pair of sharp tweezers, collected in a 24-well plate, and immersed in PBS for 20 minutes at room temperature to allow the tissue to release any residual solid stresses in the tissue (**Fig. 1B**). As a result of stress relaxation, the tissue slice becomes deformed. Then the PBS is removed and the slice is embedded in 1% agarose to hold the deformation. The tissue slice is fixed for 4 hours in 10% formalin to prevent the tissue degradation, and then was washed and stored in PBS at 4°C (**Fig. 1C**). Mice in the 22 month age group were wild type and the tissue slices were stained with DAPI (Thermo Fisher), which is a fluorescent nuclear and chromosome counterstain and can emit blue fluorescence.

#### Imaging the stress-induced deformation

With the fluorescence reporter in the tissue, the sample is imaged via Olympus FV3000 laser scanning confocal microscope using UPL SAPO10X2 (Olympus, NA 0.4, 10x magnification) air immersion objective lens (Olympus) at scanning resolutions between 512×512 pixels in FV31S-SW Viewer software (Olympus). Slices from 5–7 days and 8–12 weeks groups mice had tdTomato fluorescence and were imaged using a 561 nm excitation laser and 570–620 detection wavelength, the slices from 22 months group mice had DAPI fluorescence and were imaged using a 405 nm excitation laser and 430–470 nm detection wavelength. The tissue-agarose construct that is immersed in PBS was imaged to get the 3D structure (**Fig. 1D**).

#### Post processing of the stress-induced deformation

The 3D images (**Fig. 1I**) are exported from ImageJ to MATLAB for post-processing. The post-processing includes normalized deformation calculation, and the reconstruction of the deformation map. The release of residual solid stress via slicing results in surface bending of the slice. The overall bending extent after slicing represents the area strain, a quantitative index for residual solid stress in the slice. We defined the normalized deformation, D_n_, as the average height difference from the curved surface to the midline (**Fig. 1E**).

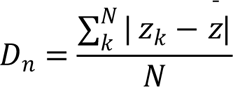

Area ratio, R_a_, is defined as the ratio of surface area, A_s_, to projection area, A_p_, represents the deformation extent (**Fig. 1F**).

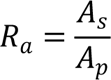

The deformation map shows the distribution of deformation in individual slices (**Fig. 1J**). Mean curvature, K_m_, is defined as the average of mean curvature on tissue slices, and indicates the degree of deformation curved (**Fig. 1G**). The curvature maps show the distribution of curvature in individual slices (**Fig. 1K**).

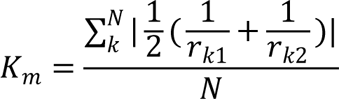

### Measurement of viscoelastic properties

#### Unconfined compression test

The bulk viscoelastic properties of the tissues were measured via the Instron 5900 Series System (Illinois Tool Works Inc) in macroscale with an unconfined compression test. Organs were sliced with the Compresstome using the slicing method with a 2 mm thickness. The slices were then punched with biopsy punches for 6 mm diameter and maintained in PBS with protease inhibitors at 4°C before mechanical testing. All measurements were performed in near-physiological PBS at room temperature. The sample was placed in a loading machine and the displacement was zeroed at the point where the upper plate was in contact with the sample, then applying a constant strain. Each specimen was compressed by 5% of original height in ramps of 1s and allowed to relax for 3 minutes. Four consecutive steps were performed to apply the displacement changing with time. The Instron then measured the force changes in the whole process, including the increase of force while compressing, and the force decreasing during the relaxation time till it reached an equilibrium point. The plots of displacement versus time and force versus time were converted to plots of stress versus time and strain versus time (**Fig. S5**). Stress modulus, σ, was the ratio of force to surface area, and strain was the ratio of displacement to total thickness. The stress is then plotted as a function of strain, and the Young’s modulus, E, is estimated as the slope of the linear fit to the stress-strain data as modulus measurement (**Fig. S5**).

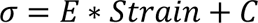

Instantaneous / equilibrium modulus ratio, R_s_, is the ratio of maximum modulus, σ_max_, to equilibrium modulus, σ_equilibrium_, which indicates how much stress the tissue releases to reach an equilibrium point (**Fig. S5**).

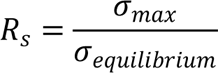

The relaxation time constant, τ, was used to compare the time of how long it takes for the tissue stress level to relax (**Fig. S5**), and can be divided into fast relaxation time constant, τ_fast_, and slow relaxation time constant, τ_slow_, for accurate evaluation.

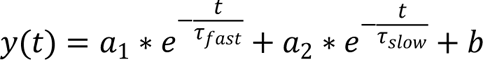

### Evaluating the effect of Ionic strength on residual solid stresses

In the slicing method, the slices are immersed in PBS for deformation due to stress relaxation after the fresh tissue been sliced. In tissue experiments, PBS is often used to maintain tissue hydration during mechanical testing, the solutes from the buffer can diffuse into the tissue and interact with its structure and mechanics (51). As the ionic strength is understood to have effect on tissue mechanics (52), the relationship between the ionic strength in PBS and the deformation on the slices is studied. Four groups of ionic strength are set through using different concentration of PBS, including 0.01X, 0.1X, 1X, and 10X PBS. Among the four groups, the 1X PBS is the physiological condition and is used in normal experiment and acts as the control group at here. The slices are immersed in specific buffer immediately after slicing for tissue deformation. Then hold the deformed slices in 1% agarose made by corresponding 0.01X, 0.1X, 1X and 10X PBS, fixed with formalin and imaged with confocal microscopy, following by tissue imaging and post-processing.

### Evaluating the spatial distribution of residual solid stress distribution in whole brain

The whole brain from mice aged 8–12 weeks was sliced to get 18 continuous coronal sections with a thickness of 250 μm each, for a total of 4500 μm. Then builds deformation and curvature maps, and quantification of normalized deformation, area ratio, and mean curvature to study the distribution of residual solid stress in brain at the coronal direction.

### Induction of Hemorrhagic Strokes

All surgical procedures were approved by the BU IACUC (Protocol number: PROTO20200045) and conducted within a designated surgical facility. All procedures were performed on C57BL/6 mTmG mice that were aged 8-12 weeks at the time of surgery under general anesthesia achieved through inhalation of isoflurane in oxygen-enriched air. Shaved mice heads were stabilized and horizontally leveled in a stereotaxic apparatus using ear bars (David Kopf, Tujunga, CA). A small craniotomy over the left coronal suture was performed using a high-speed surgical drill and visually aided by an operating microscope. A small rectangular flap of bone encompassing sections of the frontal and parietal bone was removed to expose the brain in preparation for injection. To induce hemorrhagic strokes, 1 μL of Collagenase I (Gibco™ 17018029) (0.1 U/µL in sterile PBS) was injected into the caudate putamen nucleus at 0.15 μL/min using target coordinates relative to Bregma: +1.0 mm A/P, +2.5 mm L/M and −3.0 mm D/V. A standard micropipette injection protocol was used to make all injections into the brain using pulled borosilicate glass micropipettes (WPI, Sarasota, FL, #1B100-4) that were ground to a 35° beveled tip with 150–250 μm inner diameter. Glass micropipettes were mounted to the stereotaxic frame via specialized connectors and attached, via high-pressure polyetheretherketone (PEEK) tubing, to a 10 μL syringe (Hamilton, Reno, NV, #801 RN) controlled by an automated syringe pump (Pump 11 Elite, Harvard Apparatus, Holliston, MA).

### Histology

#### Transcardial perfusions for Immunohistochemistry

After terminal anesthesia by overdose of isoflurane, mice were perfused transcardially with heparinized saline (10 units/ml of heparin) and 4% paraformaldehyde (PFA) that was prepared from 32% PFA Aqueous Solution (Cat# 15714, EMS), using a peristaltic pump at a rate of 7 mL/min. Approximately, 10 mL of heparinized saline and 50 mL of 4% PFA was used per animal. Brains were immediately dissected after perfusion and post-fixed in 4% PFA for 6-8 hours. After PFA post-fixing, brains were cryoprotected in 30% sucrose + 0.01% sodium azide in Tris-Buffered Saline (TBS) for at least 3 days with the sucrose solution replaced once after 2 days and stored at 4°C until further use.

#### Immunohistochemistry

Coronal brain sections (40 µm thick) were cut using a cryostat (Microm, HM 525). Tissue sections were stored in TBS buffer + 0.01% sodium azide at 4°C. Tissue sections were processed for immunofluorescence using free floating staining protocols described in comprehensive detail previously using donkey serum to block and triton X-100 to permeabilize tissue (53–55). The primary antibodies used are rat anti-Gfap (1:1000, Thermofisher, #13-0300); goat anti-Cd13 (1:500, R&D Systems, AF2335); guinea pig anti-NeuN (1:500, Synaptic Systems, #266 004). All secondary antibodies purchased from Jackson ImmunoResearch Laboratories, with donkey host and target specified by the primary antibody. Secondary antibodies were diluted 1:250 prior to incubation. Cell nuclei were stained with 4’,6’-diamidino-2-phenylindole dihydrochloride (DAPI; 2 ng/ml; Molecular Probes). Stained sections were imaged using epifluorescence and deconvolution epifluorescence microscopy on an Olympus IX83. Subsequently, images were prepared for publication using NIH Image J (1.53) software.

### Statistical analyses

The data are presented as mean ± standard error of the mean. Statistics between every two groups of data are calculated as two-tailed t-test (n≥3) to determine significance or p-values. Statistics among multiple data sets are calculated as one-way ANOVA test to determine significance or p-values. All p-values are reported on figures to observe statistical trends of data.

## Acknowledgments

The research reported in this publication was supported by the Boston University Micro and Nano Imaging Facility and the Office of the Director, National Institutes of Health of the National Institutes of Health. The 22 months mice were provided from Dr. Brianne Connizzo’s lab. The content is solely the responsibility of the authors and does not necessarily represent the official views of the National Institute of Health.

## Supporting Information

### Supporting Information Text

The supplementary experiments are used to test the validation of our slicing method and evaluate the accuracy of measurement and quantification. Additionally, we also evaluate the potential relationship between residual solid stress changes in the mouse brain and the volume changes in the human brain.

## Methods

### Positive and negative control organs

The heart is studied previously as an organ which has a strong residual stress in it (8). And can be used to test the deformation quantification accuracy of showing the extent of residual solid stress. The same slicing method is used to slice the mouse heart, compared the deformation and curvature maps and quantification of normalized deformation, area ratio, and mean curvature between kidney and heart.

### Evaluating the buoyancy effect

Since the tissue slice was immersed in PBS for the deformation occur, flipping test is done to avoid the possibility that the deformation was an artifact caused by the buoyancy. The buoyancy might have different pressure acting on opposite sides of the slice, and thus the heavy part will sink and the light part will go up and floating in the liquid, and result to deformation. Imaging the slice immediately after tissue deformed, and then flip the slice to upside down and then image again. These all steps will be finished before the tissue degradation. Then compared the cross-section view and the quantification of normalized deformation, area ratio, and mean curvature.

### Evaluating the effect of fixation

In the slicing method, formalin is used to fix the tissue and avoid tissue degradation, the potential artifact of fixation needs to be evaluated. The brain slice is imaged after it deformed and hold in 1% agarose, and then do the fixation with formalin, wash with PBS, after that did another imaging, and compare the cross-section view, deformation and curvature maps, and quantification of normalized deformation, area ratio, and mean curvature.

### Fitting the changing tendency between residual solid stress and brain volume

The mouse age is converted to human age as showed in previous study (56), that 5–7 day mice are equivalent to 0.1 human years, 8-12 week mice are equivalent to 20 human years, and 22 month mice are equivalent to 80 human years. The changing tendencies of normalized deformation, area ratio, and mean curvature in mouse brain with respect to age closely trend with changes in brain volume as humans age (26, 57, 58).

## Author Contributions

S.Zheng and H.T.N designed the study; S.Zheng, S.Zhang, and E.D performed research; S.Zheng, R.B, R.L, and H.T.N contributed to data analysis; S.Zheng analyzed data; S.Zheng, R.B, R.L, S.Zhang, E.D, T. O., and H.T.N wrote the manuscript.

## Competing Interest Statement

Authors declare that they have no competing interests.

## Data and materials availability

All data are available in the main text or the supplementary materials. Raw data and code for analysis are available upon request.

## Classification

Biological Sciences, Biophysics and Computational Biology

---

**Fig. S1.**
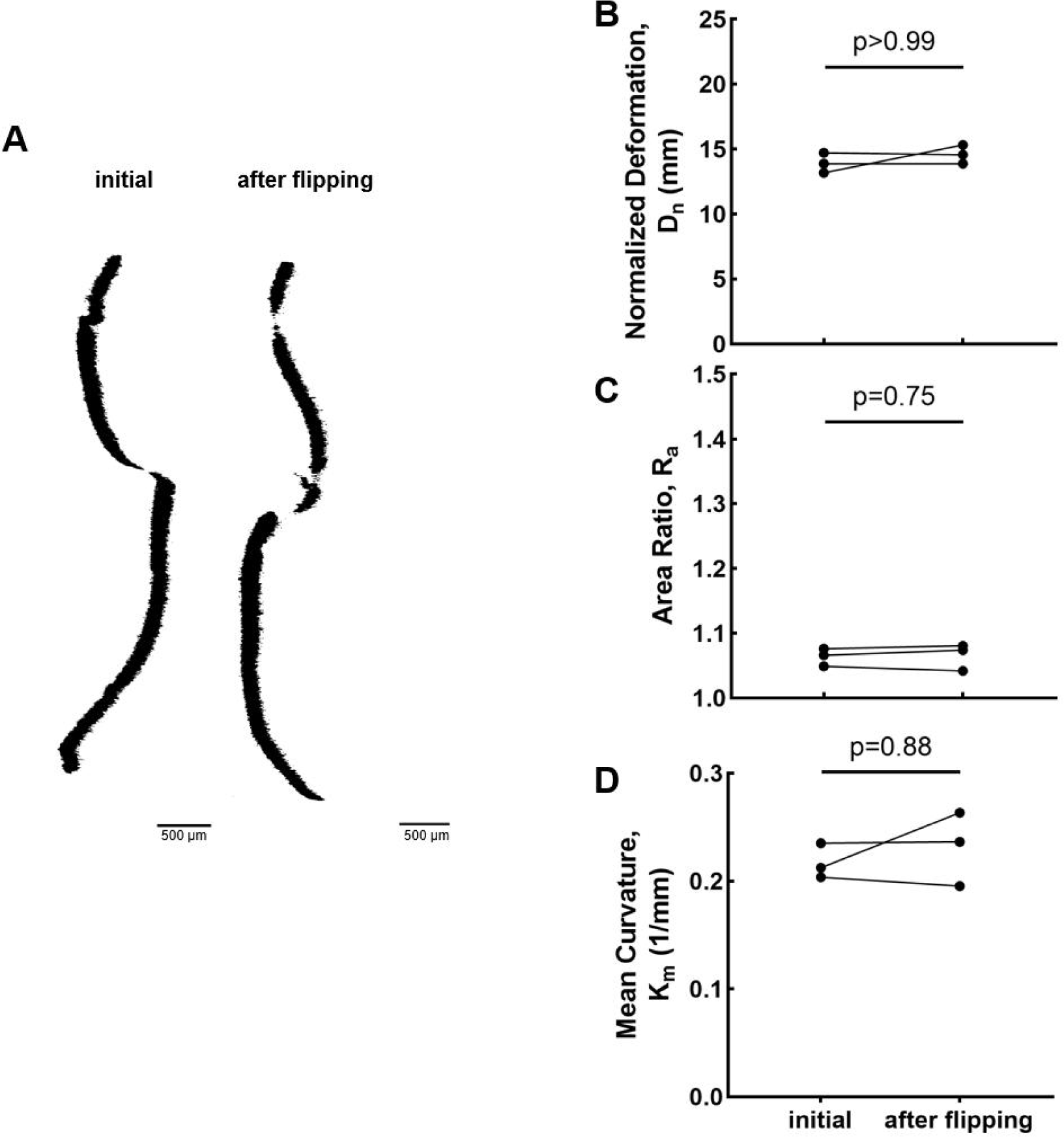
Buoyancy does not affect residual solid stress quantification. (**A**) The orthogonal view of representative microscopy images of tissue slices before and after flipping. (**B**) Statistics of normalized deformation, (**C**) area ratio, and (**D**) mean curvature between slices initial and slices after flipping (mean ± SEM, n=3 slices, two-tailed t-test). All normalized deformation, area ratio, and mean curvature have no significant difference of brain slices between initial and after flipping stages.

**Fig. S2.**
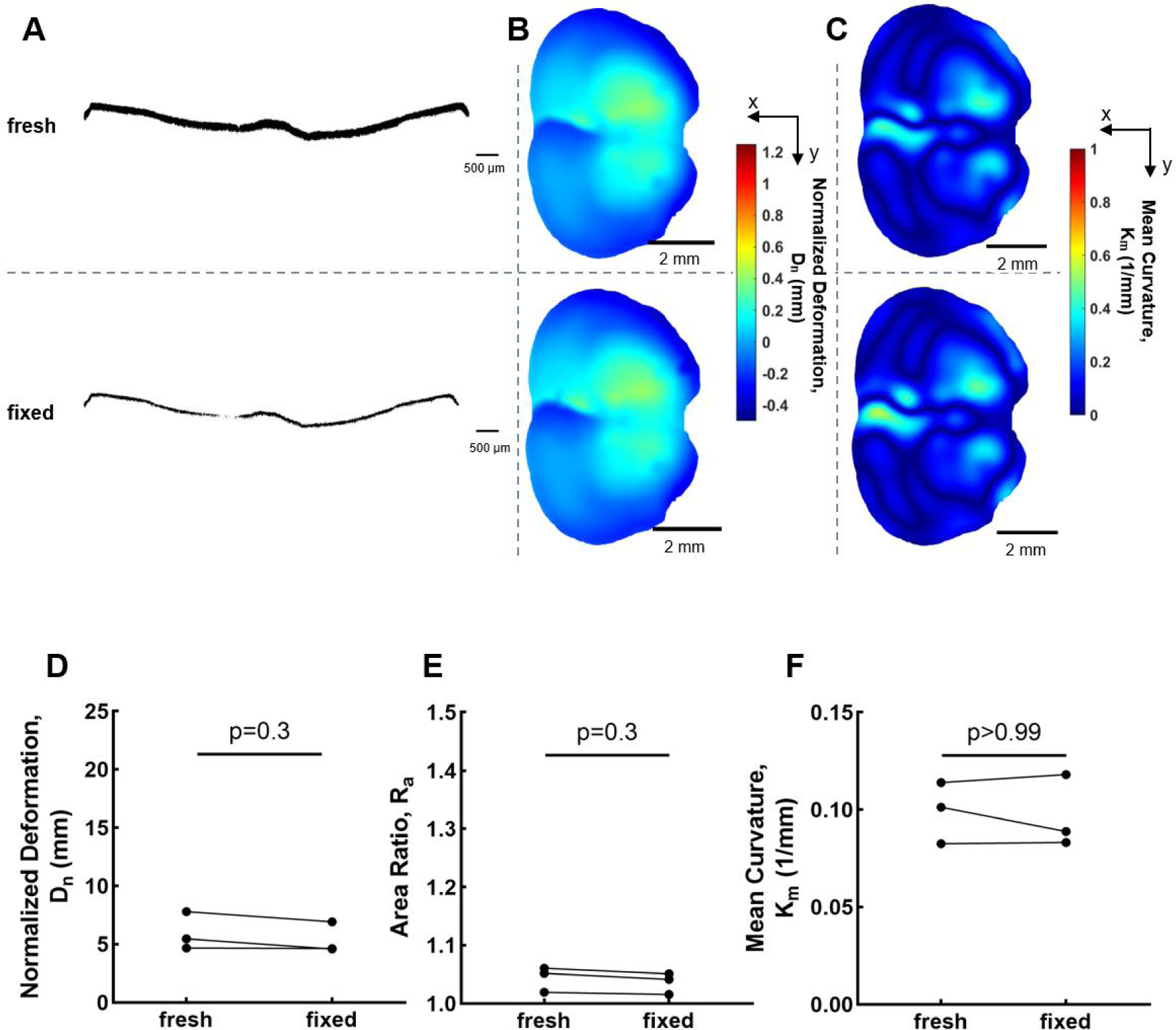
Fixation does not affect residual solid stress quantification. (**A**) The orthogonal view of microscopy images of the tissue slices, (**B**) deformation maps, and (**C**) mean curvature maps of representative brain slices from fresh and fixed states. Statistics of (**D**) normalized deformation, (**E**) area ratio, and (**F**) mean curvature between fresh and fixed brain slices (mean ± SEM, n=3 slices, two-tailed t-test). All normalized deformation, area ratio, and mean curvature have no significant difference of brain slices between fresh and fixed stages.

**Fig. S3.**
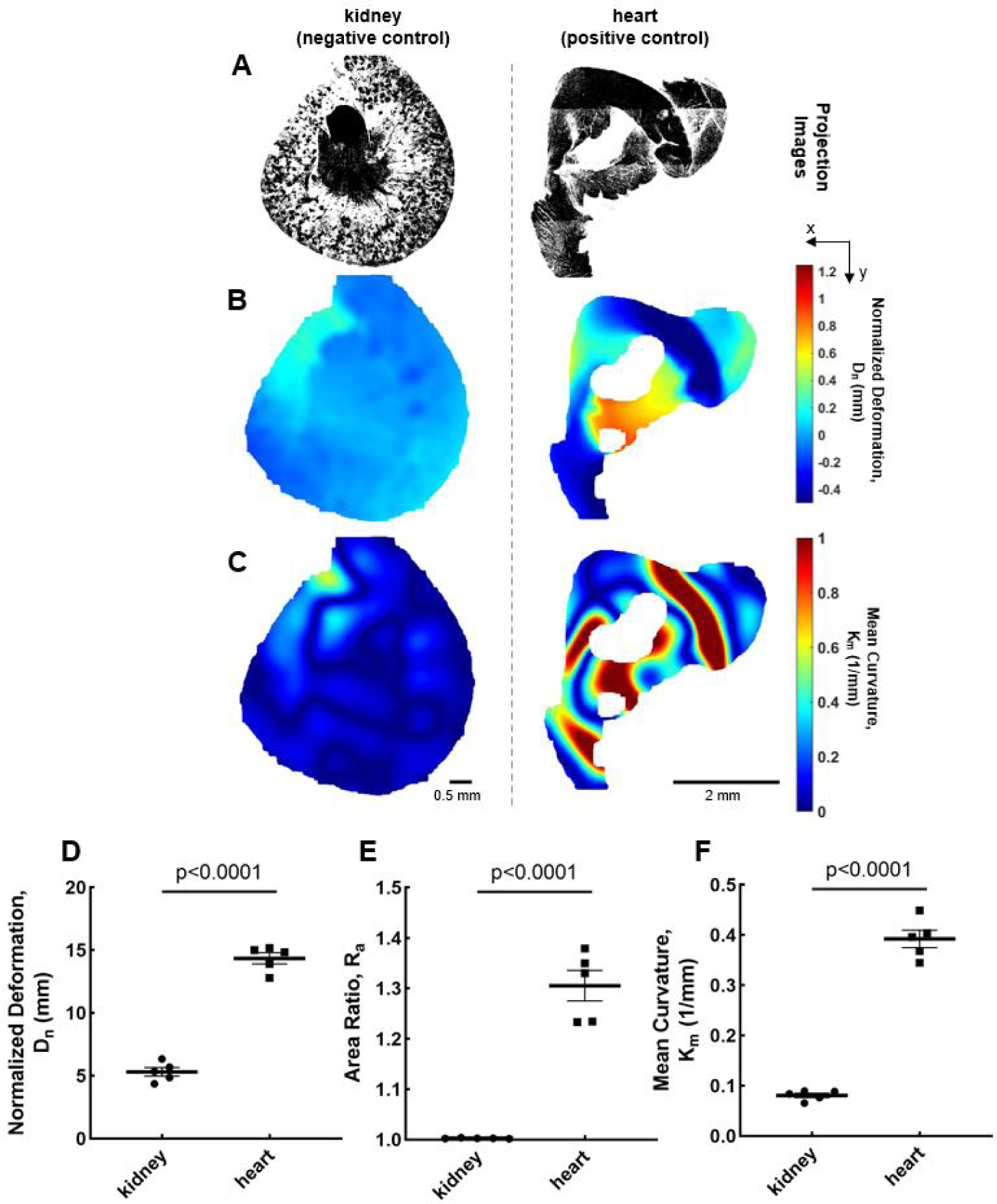
Higher residual solid stress exists in heart. (**A**) Projected microscopy images, (**B**) corresponding deformation maps, and (**C**) mean curvature maps of representative slices from kidney and heart. Statistics of (**D**) normalized deformation, (**E**) area ratio, and (**F**) mean curvature among kidney and heart slices (mean ± SEM, N=5 mice, two-tailed t-test).

**Fig. S4.**
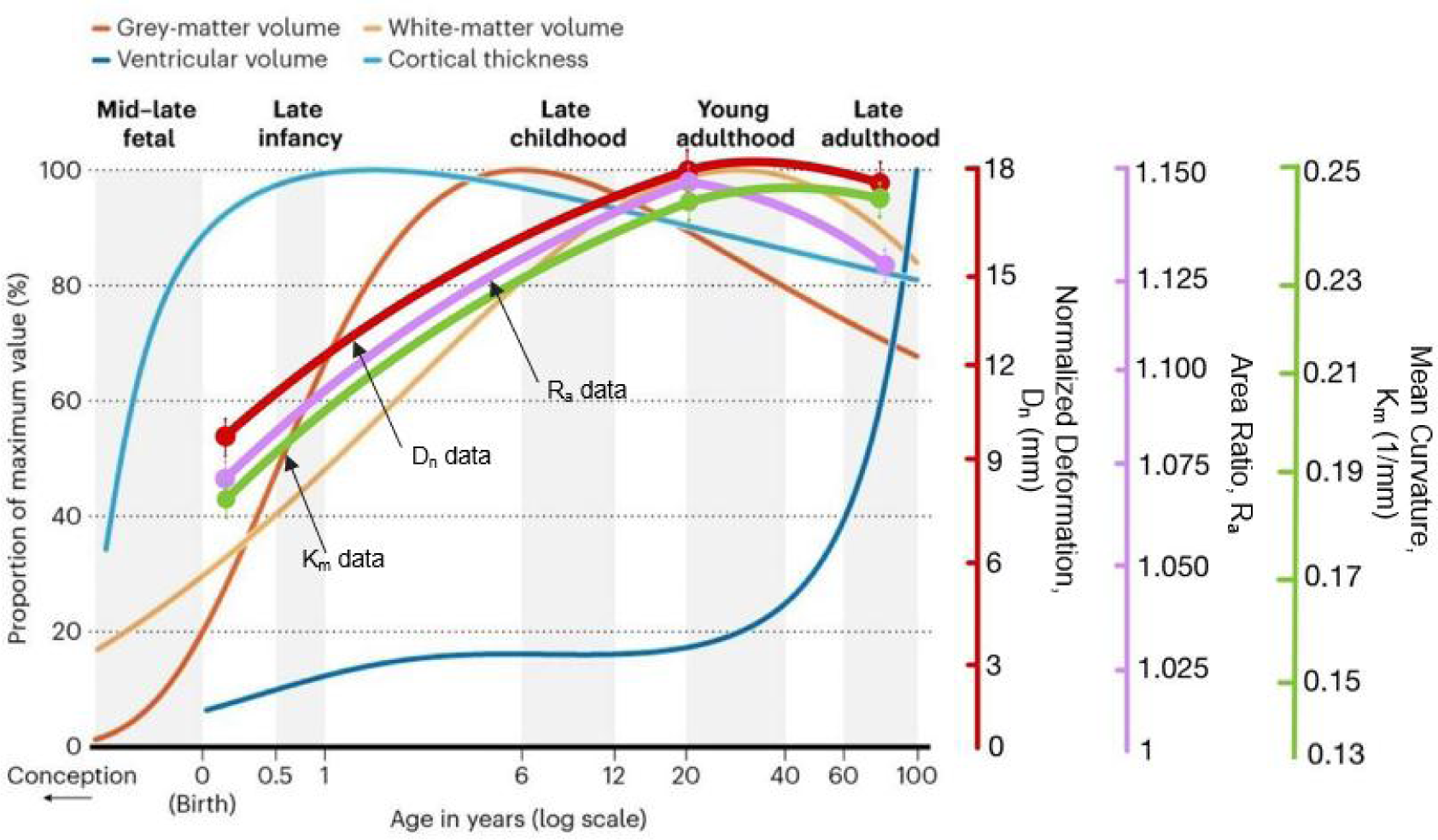
The comparison between volume change in human brain and residual solid stress change in mouse brain. Convert mouse lifespan to human (56) where 5–7 day, 8-12 week, and 22 month mice are equivalent to 0.1, 20, and 80 years in human, respectively. The normalized deformation, D_n_, area ratio, R_a_, and mean curvature, K_m_, trend with age in the brain compared to the volume changes. Modified from (26, 57, 58).

**Fig. S5.**
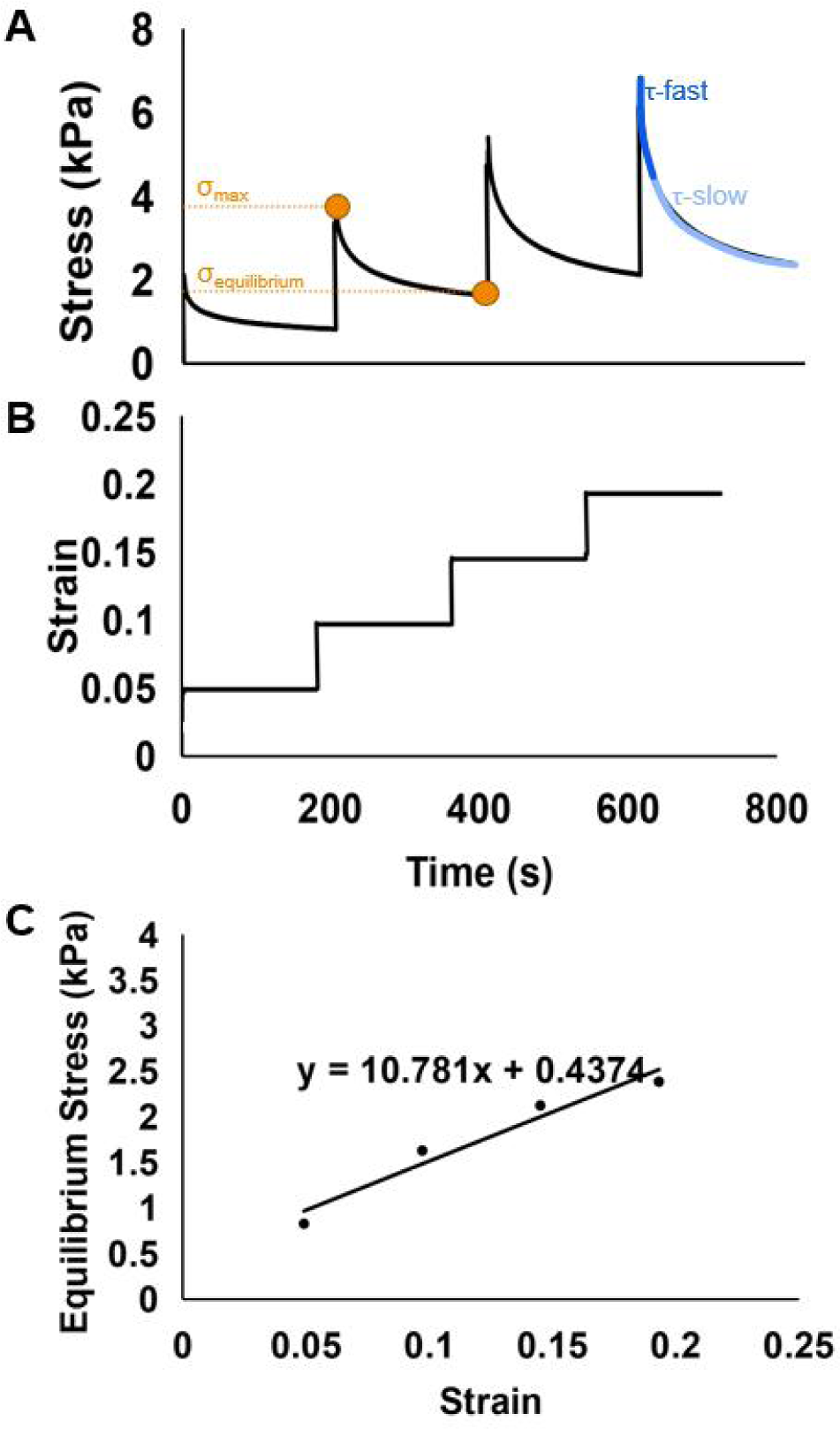
Methodology for quantification of viscoelastic properties. (A) The stress relaxation is measured with an unconfined compression test on an Instron. From the curves, the instantaneous / equilibrium modulus ratio, R_s_, is the ratio of maximum, σ_max_, and equilibrium modulus, σ_equilibrium_ in each step, indicating how much stress the tissue released to reach a lower equilibrium point. Relaxation time constant, τ, evaluates the time of how long it takes for the tissue stress level to relax and become stable, and can be divided into fast, τ_fast_, and slow parts, τ_slow_. (**B**) Four steps each of 5% compressive is applied and held for 3 minutes to let the stress relax. (**C**) The equilibrium stress is plotted as a function of strain and the Young’s modulus is estimated as the slope of the linear fit to the stress-strain data.

## Notes

### Competing Interest Statement

The authors have declared no competing interest.

## References

1. H. T. Nia, L. L. Munn, R. K. Jain, Physical traits of cancer. Science 370 (2020).

2. M. Kalli, T. Stylianopoulos, Defining the Role of Solid Stress and Matrix Stiffness in Cancer Cell Proliferation and Metastasis. Front Oncol 8, 55 (2018).

3. T. R. Huycke et al., Genetic and Mechanical Regulation of Intestinal Smooth Muscle Development. Cell 179, 90–105 e121 (2019).

4. M. Pesce et al., Cardiac fibroblasts and mechanosensation in heart development, health and disease. Nat Rev Cardiol 20, 309–324 (2023).

5. H. I. Harn et al., The tension biology of wound healing. Exp Dermatol 28, 464–471 (2019).

6. Y. C. Fung, S. Q. Liu, Change of residual strains in arteries due to hypertrophy caused by aortic constriction. Circ Res 65, 1340–1349 (1989).

7. Z. Chen, Q. Guo, E. Dai, N. Forsch, L. A. Taber, How the embryonic chick brain twists. J R Soc Interface 13 (2016).

8. L. A. Taber, Biomechanics of cardiovascular development. Annu Rev Biomed Eng 3, 1–25 (2001).

9. G. Helmlinger, P. A. Netti, H. C. Lichtenbeld, R. J. Melder, R. K. Jain, Solid stress inhibits the growth of multicellular tumor spheroids. Nat Biotechnol 15, 778–783 (1997).

10. H. T. Nia, et al., Solid stress and elastic energy as measures of tumour mechanopathology. Nat Biomed Eng 1 (2016).

11. T. Stylianopoulos et al., Causes, consequences, and remedies for growth-induced solid stress in murine and human tumors. Proc Natl Acad Sci U S A 109, 15101–15108 (2012).

12. M. E. Fernandez-Sanchez et al., Mechanical induction of the tumorigenic beta-catenin pathway by tumour growth pressure. Nature 523, 92–95 (2015).

13. P. M. Munne et al., Compressive stress-mediated p38 activation required for ERalpha + phenotype in breast cancer. Nat Commun 12, 6967 (2021).

14. D. Jones et al., Solid stress impairs lymphocyte infiltration into lymph-node metastases. Nat Biomed Eng 5, 1426–1436 (2021).

15. V. P. Chauhan et al., Angiotensin inhibition enhances drug delivery and potentiates chemotherapy by decompressing tumour blood vessels. Nat Commun 4, 2516 (2013).

16. T. P. Padera et al., Pathology: cancer cells compress intratumour vessels. Nature 427, 695 (2004).

17. J. M. Tse et al., Mechanical compression drives cancer cells toward invasive phenotype. Proc Natl Acad Sci U S A 109, 911–916 (2012).

18. G. Seano et al., Solid stress in brain tumours causes neuronal loss and neurological dysfunction and can be reversed by lithium. Nat Biomed Eng 3, 230–245 (2019).

19. H. T. Nia et al., In vivo compression and imaging in mouse brain to measure the effects of solid stress. Nat Protoc 15, 2321–2340 (2020).

20. E. Moeendarbary et al., The soft mechanical signature of glial scars in the central nervous system. Nat Commun 8, 14787 (2017).

21. A. Goriely et al., Mechanics of the brain: perspectives, challenges, and opportunities. Biomech Model Mechanobiol 14, 931–965 (2015).

22. C. M. Hall, E. Moeendarbary, G. K. Sheridan, Mechanobiology of the brain in ageing and Alzheimer’s disease. Eur J Neurosci 53, 3851–3878 (2021).

23. S. Budday et al., Mechanical properties of gray and white matter brain tissue by indentation. J Mech Behav Biomed Mater 46, 318–330 (2015).

24. G. Xu, P. V. Bayly, L. A. Taber, Residual stress in the adult mouse brain. Biomech Model Mechanobiol 8, 253–262 (2009).

25. T. Tuomas et al., On the growth and form of cortical convolutions. Nature Physics 12, 588–593 (2016).

26. R. A. I. Bethlehem et al., Brain charts for the human lifespan. Nature 604, 525–533 (2022).

27. A. Vipin et al., Cerebrovascular disease influences functional and structural network connectivity in patients with amnestic mild cognitive impairment and Alzheimer’s disease. Alzheimers Res Ther 10, 82 (2018).

28. J. M. Barnes, L. Przybyla, V. M. Weaver, Tissue mechanics regulate brain development, homeostasis and disease. J Cell Sci 130, 71–82 (2017).

29. L. Puy et al., Intracerebral haemorrhage. Nat Rev Dis Primers 9, 14 (2023).

30. R. W. Regenhardt, A. S. Das, E. H. Lo, L. R. Caplan, Advances in Understanding the Pathophysiology of Lacunar Stroke: A Review. JAMA Neurol 75, 1273–1281 (2018).

31. S. Narasimhan et al., Development of a mechanics-based model of brain deformations during intracerebral hemorrhage evacuation, SPIE Medical Imaging (SPIE, 2017), vol. 10135.

32. H. T. Nia et al., Quantifying solid stress and elastic energy from excised or in situ tumors. Nat Protoc 13, 1091–1105 (2018).

33. Z. Sue et al., In vivo multiscale measurements of solid stresses in tumors reveal scale-dependent stress transmission. Nat Biomed Eng 10.21203/rs.3.rs-1697924/v1 (2023).

34. M. E. Dolega et al., Cell-like pressure sensors reveal increase of mechanical stress towards the core of multicellular spheroids under compression. Nat Commun 8, 14056 (2017).

35. W. Lee et al., Dispersible hydrogel force sensors reveal patterns of solid mechanical stress in multicellular spheroid cultures. Nat Commun 10, 144 (2019).

36. E. Mohagheghian et al., Quantifying compressive forces between living cell layers and within tissues using elastic round microgels. Nat Commun 9, 1878 (2018).

37. O. Campas et al., Quantifying cell-generated mechanical forces within living embryonic tissues. Nat Methods 11, 183–189 (2014).

38. O. Chaudhuri, J. Cooper-White, P. A. Janmey, D. J. Mooney, V. B. Shenoy, Effects of extracellular matrix viscoelasticity on cellular behaviour. Nature 584, 535–546 (2020).

39. A. J. Grodzinsky, E. H. Frank, Fields, forces, and flows in biological systems (Garland Science, London; New York, 2011), pp. xii, 308 p.

40. S. Dauth et al., Extracellular matrix protein expression is brain region dependent. J Comp Neurol 524, 1309–1336 (2016).

41. C. Voutouri, C. Polydorou, P. Papageorgis, V. Gkretsi, T. Stylianopoulos, Hyaluronan-Derived Swelling of Solid Tumors, the Contribution of Collagen and Cancer Cells, and Implications for Cancer Therapy. Neoplasia 18, 732–741 (2016).

42. S. R. Eisenberg, A. J. Grodzinsky, Swelling of articular cartilage and other connective tissues: electromechanochemical forces. J Orthop Res 3, 148–159 (1985).

43. B. W. Bigger, D. J. Begley, D. Virgintino, A. V. Pshezhetsky, Anatomical changes and pathophysiology of the brain in mucopolysaccharidosis disorders. Mol Genet Metab 125, 322–331 (2018).

44. N. B. Schwartz, M. S. Domowicz, Proteoglycans in brain development and pathogenesis. FEBS Lett 592, 3791–3805 (2018).

45. D. I. Zafeiriou, S. P. Batzios, Brain and spinal MR imaging findings in mucopolysaccharidoses: a review. AJNR Am J Neuroradiol 34, 5–13 (2013).

46. N. B. Schwartz, M. S. Domowicz, Chemistry and Function of Glycosaminoglycans in the Nervous System. Adv Neurobiol 29, 117–162 (2023).

47. A. I. Qureshi et al., Spontaneous intracerebral hemorrhage. N Engl J Med 344, 1450–1460 (2001).

48. J. A. Caceres, J. N. Goldstein, Intracranial hemorrhage. Emerg Med Clin North Am 30, 771–794 (2012).

49. G. R. Sutherland, R. N. Auer, Primary intracerebral hemorrhage. J Clin Neurosci 13, 511–517 (2006).

50. H. Zhu et al., Role and mechanisms of cytokines in the secondary brain injury after intracerebral hemorrhage. Prog Neurobiol 178, 101610 (2019).

51. B. N. Safa, K. D. Meadows, S. E. Szczesny, D. M. Elliott, Exposure to buffer solution alters tendon hydration and mechanics. J Biomech 61, 18–25 (2017).

52. A. M. Nguyen, M. E. Levenston, Comparison of osmotic swelling influences on meniscal fibrocartilage and articular cartilage tissue mechanics in compression and shear. J Orthop Res 30, 95–102 (2012).

53. T. M. O’Shea et al., Foreign body responses in mouse central nervous system mimic natural wound responses and alter biomaterial functions. Nat Commun 11, 6203 (2020).

54. M. A. Anderson et al., Required growth facilitators propel axon regeneration across complete spinal cord injury. Nature 561, 396–400 (2018).

55. T. M. O’Shea et al., Lesion environments direct transplanted neural progenitors towards a wound repair astroglial phenotype in mice. Nat Commun 13, 5702 (2022).

56. S. Dutta, P. Sengupta, Men and mice: Relating their ages. Life Sci 152, 244–248 (2016).

57. M. Kozlov, Your brain expands and shrinks over time - these charts show how. Nature 604, 230–231 (2022).

58. S. Marek et al., Reproducible brain-wide association studies require thousands of individuals. Nature 603, 654–660 (2022).

