## Supplementary Figures for "Alteration of mechanical stresses in the murine brain by age and hemorrhagic stroke"

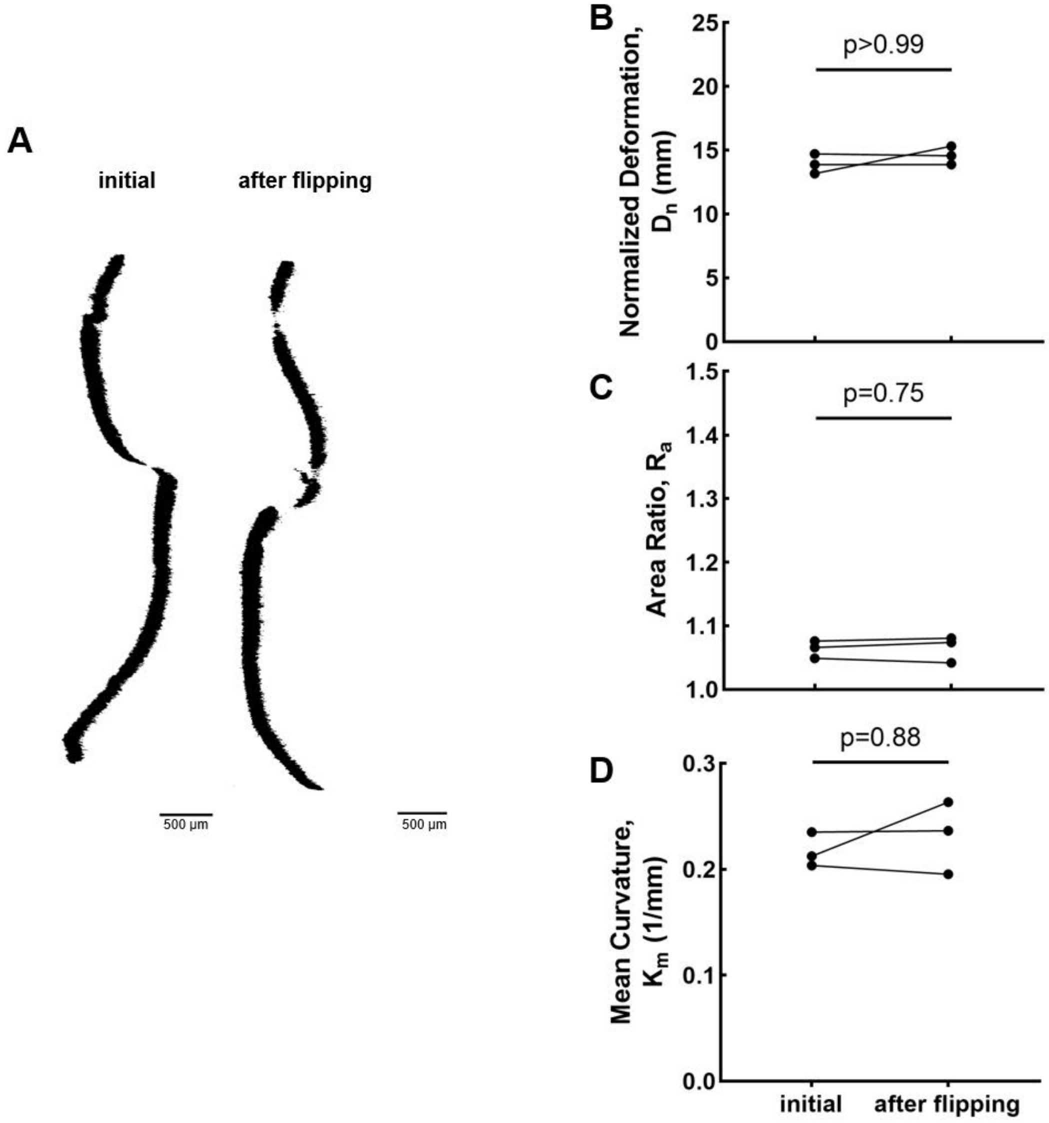


**Fig. S1. Buoyancy does not affect residual solid stress quantification.** (**A**) The orthogonal view of representative microscopy images of tissue slices before and after flipping. (**B**) Statistics of normalized deformation, (**C**) area ratio, and (**D**) mean curvature between slices initial and slices after flipping (mean ± SEM, n=3 slices, two-tailed t-test). All normalized deformation, area ratio, and mean curvature have no significant difference of brain slices between initial and after flipping stages.

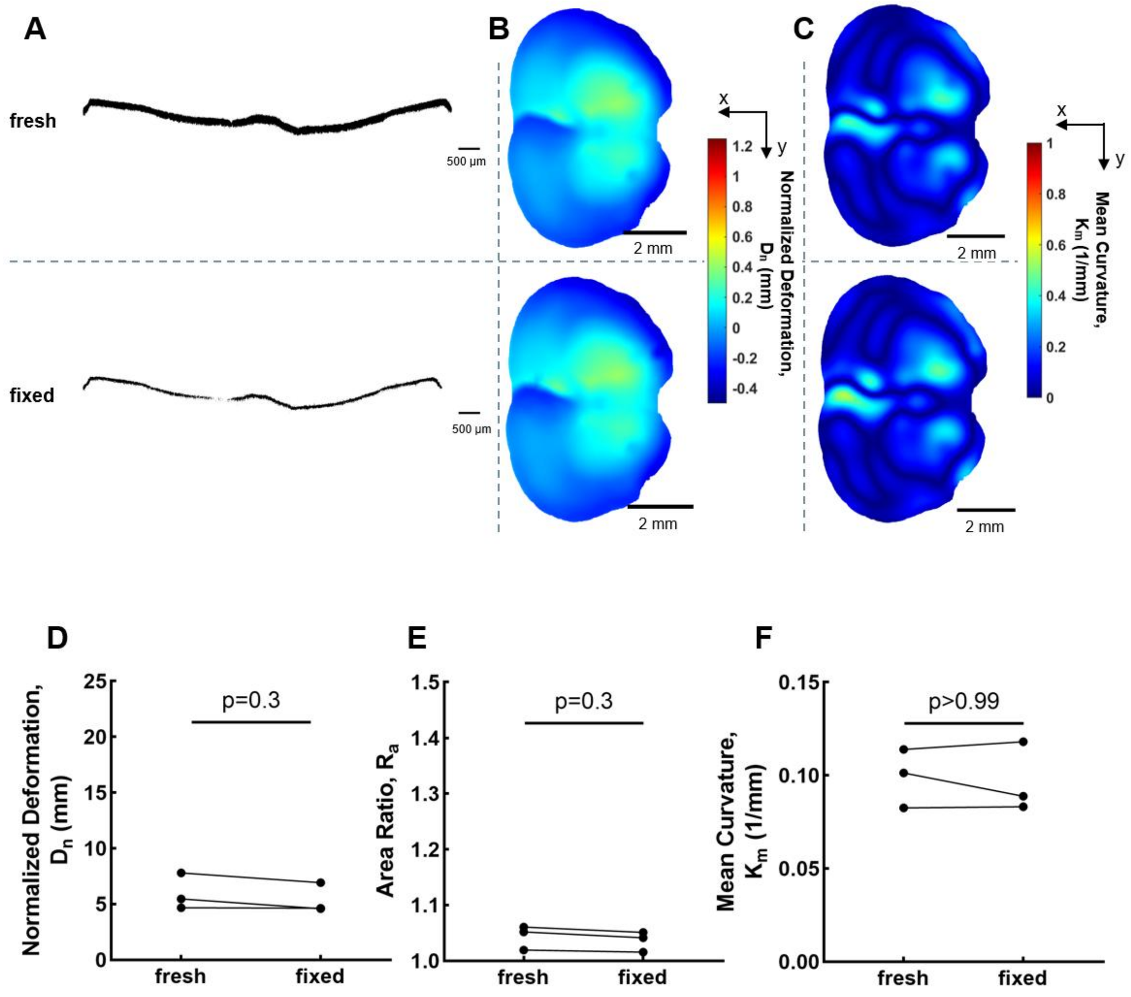


**Fig. S2. Fixation does not affect residual solid stress quantification.** (**A**) The orthogonal view of microscopy images of the tissue slices, (**B**) deformation maps, and (**C**) mean curvature maps of representative brain slices from fresh and fixed states. Statistics of (**D**) normalized deformation, (**E**) area ratio, and (**F**) mean curvature between fresh and fixed brain slices (mean ± SEM, n=3 slices, two-tailed t-test). All normalized deformation, area ratio, and mean curvature have no significant difference of brain slices between fresh and fixed stages.

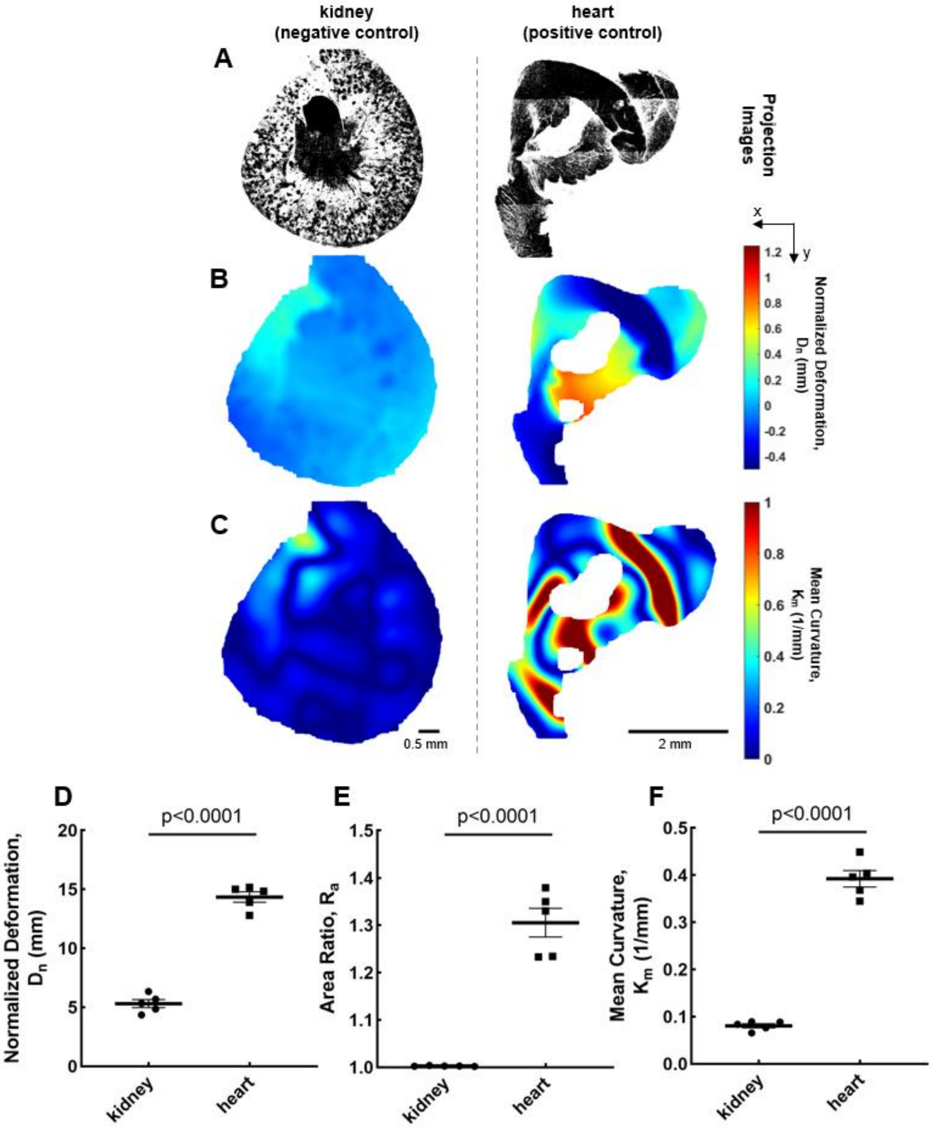


**Fig. S3. Higher residual solid stress exists in heart.** (**A**) Projected microscopy images, (**B**) corresponding deformation maps, and (**C**) mean curvature maps of representative slices from kidney and heart. Statistics of (**D**) normalized deformation, (**E**) area ratio, and (**F**) mean curvature among kidney and heart slices (mean ± SEM, N=5 mice, two-tailed t-test).

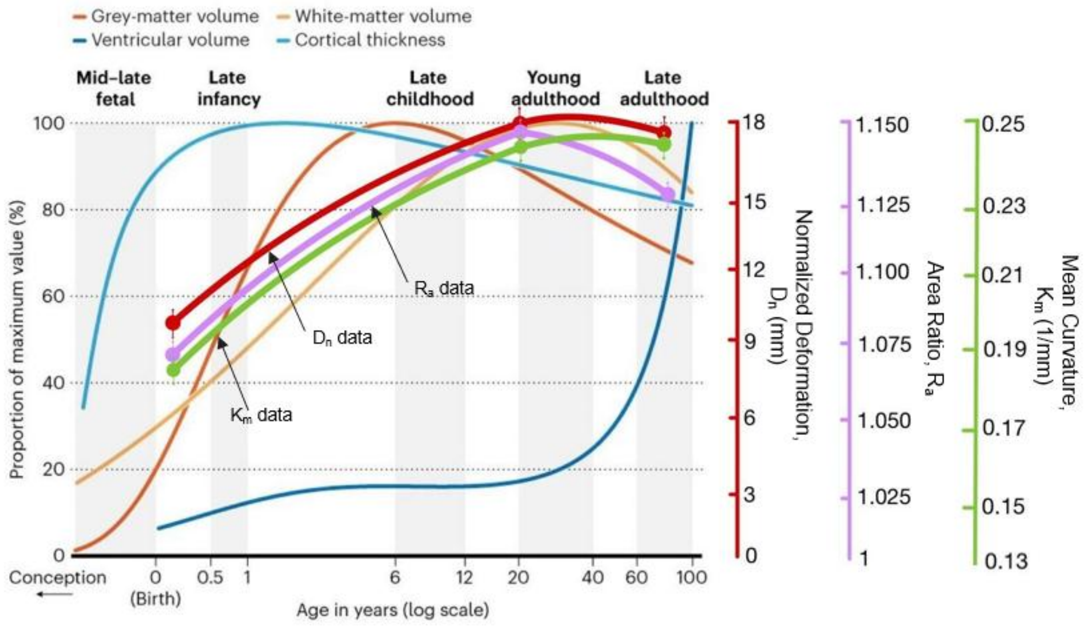


**Fig. S4. The comparison between volume change in human brain and residual solid stress change in mouse brain.** Convert mouse lifespan to human (56) where 5–7 day, 8-12 week, and 22 month mice are equivalent to 0.1, 20, and 80 years in human, respectively. The normalized deformation, $D_n$, area ratio, $R_a$, and mean curvature, $K_m$, trend with age in the brain compared to the volume changes. Modified from (26, 57, 58).

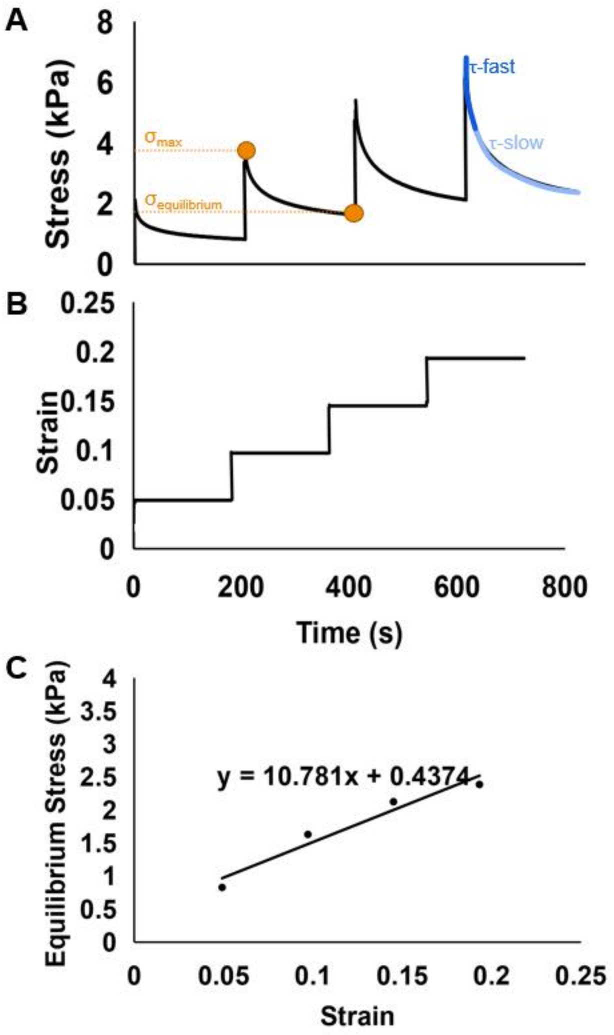


**Fig. S5. Methodology for quantification of viscoelastic properties. (A)** The stress relaxation is measured with an unconfined compression test on an Instron. From the curves, the instantaneous / equilibrium modulus ratio, $R_s$, is the ratio of maximum, $\sigma_{max}$, and equilibrium modulus, $\sigma_{equilibrium}$ in each step, indicating how much stress the tissue released to reach a lower equilibrium point. Relaxation time constant, τ, evaluates the time of how long it takes for the tissue stress level to relax and become stable, and can be divided into fast, $\tau_{fast}$, and slow parts, $\tau_{slow}$. (**B**) Four steps each of 5% compressive is applied and held for 3 minutes to let the stress relax. (**C**) The equilibrium stress is plotted as a function of strain and the Young's modulus is estimated as the slope of the linear fit to the stress-strain data.
